# Cooperation, only for high rewards – a solvable task-based study on free-ranging dogs

**DOI:** 10.1101/736009

**Authors:** Debottam Bhattacharjee, Rohan Sarkar, Shubhra Sau, Daisy Babu, Asawari Albal, Diksha Mehta, Anindita Bhadra

## Abstract

The benefits of group living mostly surpass the disadvantages like sharing of resources and competition over food, space and mates, driving the evolution of social organization. Group living can be facilitated by social tolerance and cooperation among the group members. Social canids (e.g. wolves) display cooperative breeding, hunting, and prosocial activities in different contexts. Unlike cooperative pack-living wolves (*Canis lupus lupus*), their descendants, domesticated dogs (*Canis lupus familiaris*), show varying levels of associations from solitary to stable social groups. Free-ranging dogs are group-living but prefer to forage solitarily, hence providing an excellent opportunity for investigating social tolerance and coordinated task performance among the members in various situations. We tested 113 adult-only groups of free-ranging dogs in three different tasks to investigate group responses and performance in problem-solving situations in the presence of an unfamiliar human. Task 1 (unfamiliar, single food reward) and 2 (familiar, single food reward) examined group responses and cooperation from the perspective of familiarity, while Task 3 (familiar, multiple food rewards) enabled us to test whether increased food rewards promote social tolerance and food sharing among the group members. Regardless of significantly higher performance in Task 2 compared to Task 1, cooperation and food sharing were significantly lower in both. Task 3 revealed a strong positive correlation between food sharing and social tolerance, but not between success and social tolerance, suggesting a tendency for cooperation. We conclude that context-dependent cooperation and tolerance among group members facilitate group-living in free-ranging dogs.

**Significance statement:** Group living is a common phenomenon in the animal world where the members of a group show social tolerance and co-operative behaviours towards each other. This need for cooperative intents increases manifolds while groups face different problem-solving situations in their day to day lives. Here, we tested a large number of free-ranging dog groups to understand general cooperative intents such as social tolerance and food sharing in different problem-solving conditions. We found shreds of evidence of context-dependent cooperation and social tolerance among group members with minimal display of aggression. It is not adaptive for the dogs to fight or display aggression over resources. Alternatively, use of subtle cues such as display of dominance and subordination seem to be more plausible mechanisms for the development of efficient scavenging strategies and maintaining hierarchy.

## Introduction

A wide range of species display differing levels of social organization, from loose groups like herds, to highly organized societies like in the social insects. Group-living requires cooperation among individuals (Buss 1981; McCallum et al. 1985) and simultaneous or co-ordinated actions over varied tasks like foraging (Clark and Mangel 1986), hunting (Packer and Ruttan 1988; Stander 1992; Creel 1997), protection of nests (Lazaro-Perea 2001; Schradin 2004; Brown 2013), rearing of offspring (Stacey and Ligon 1991; Clutton-Brock 2002), etc. Social behaviour has evolved as an evolutionarily stable strategy across taxa, through multiple selection events, as the advantages of living in groups compensates for the obvious disadvantages involved in the process, like the sharing of resources (Axelrod and Hamilton 1981; Kapheim et al. 2015). Sociality involves the emergence of coordination among members and subsequent cooperation through resolution of conflict, and is thus a dynamic process (Monnin and Ratnieks 1999; Connor 2000; Franz et al. 2013). Intragroup cooperation sometime helps to enhance the fitness of the members through increased reproduction, while in some cases, cooperation is imperative for survival in a harsh environment (Gittleman 1989; Ebensperger et al. 2012). Cooperation has been suggested to correlate with high social tolerance and low aggression towards group members (Werdenich and Huber 2002; Scott 2006). Thus, studying basic components of cooperation, like food sharing, social tolerance, allo-parenting etc. can help to develop an understanding of the evolution of group dynamics in species.

Canids display a wide diversity of social organization from large groups or packs found in species like wolves (Fox 1971; Macdonald 1983), dholes (Macdonald 1983), etc. to species that live in small groups like foxes (Fox 1971; Lloyd 1981) and jackals (Macdonald 1983). Descendants of the gray wolves, the domestic dogs (*Canis lupus familiaris*) are an interesting example of canids that can live as pets and also in social groups with interesting social dynamics, as free-ranging populations (Sen Majumder et al., 2014; Paul and Bhadra 2018). Though most of our current understanding of the behaviour, cognitive abilities and evolutionary history of dogs is based on studies with pets, majority of the world’s dog population is actually free-ranging (Lord et al. 2013), localized mostly in developing nations. They live without direct human supervision in human-dominated habitats (Cafazzo, Valsecchi, Bonanni & Natoli, 2010; Hughes & Macdonald, 2013; Sen Majumder et al., 2014a; Vanak & Gompper, 2009). Several studies have been carried out with individual free-ranging dogs to understand their physical and social cognitive abilities, in contexts like food preference (Bhadra et al. 2016), task-solving (Bhattacharjee, Dasgupta, et al., 2017; Brubaker, Dasgupta, Bhattacharjee, Bhadra, & Udell, 2017) and interspecific association with humans (Bhattacharjee et al. 2017b, c), but similar studies have not been conducted with groups of free-ranging dogs.

Free-ranging dogs live in social groups of varying sizes (2 - 15 individuals, from observations). As scavengers, they forage solitarily most of the time, though this tendency can change during seasons like mating, pup-emergence, etc., when group foraging increases (Sen Majumder et al., 2014). They are known to scavenge together over large and open garbage dumps mostly without conflict and aggression (Bhadra et al., 2016; Sen Majumder et al., 2014). In free-ranging dog groups, mothers provide extensive care to their pups, but also display conflict over food sharing during the weaning period (Paul & Bhadra, 2017; Paul, Sen Majumder & Bhadra, 2014b). Allo-parental care is often observed to be provided by both females and males within groups (Paul, Sen Majumder, & Bhadra, 2014a). Thus, free-ranging dog groups show interesting cooperation-conflict dynamics in contexts of parental care and foraging.

It has previously been shown that wolves better cooperate with their pack members in a string-pulling task compared to similarly kept and raised dogs (Marshall-Pescini et al. 2017). Pack-living dogs have been shown to share food with members based on rank positioning, suggesting a role of dominance hierarchy (Dale et al. 2017). Moreover, a steeper dominance hierarchy in such dogs compared to similarly raised wolves has also been reported (Range et al. 2015). Unfortunately, studies are greatly lacking pertaining to free-ranging dogs’ group performance and cooperation in problem-solving situations, which could give us insights into the maintenance of group cohesiveness and social hierarchy. Individual free-ranging dogs have been shown to depend on humans when faced with an unfamiliar task, exhibiting proximity-seeking and gazing behaviours (Bhattacharjee et al. 2017a). While it is essential to test their behaviours individually, it is also necessary to investigate the group responses to check if the dogs seek help from group members in similar situations and if members of a group help each other to solve a task and share food. We carried out field-based experiments with free-ranging dog groups to test their responses in an unfamiliar (Task 1) and two familiar tasks (Task 2 and Task 3) with different amounts of food rewards in the presence of an unfamiliar human experimenter. Tasks 1 and 2 provided an option of a moderately large piece of raw chicken as a food reward, while Task 3 provided a considerably higher amount of food reward in a familiar set-up. In Tasks 1 and 2, we checked how familiarity influences the problem-solving ability of dogs when present in groups. Task 3 differed from the other two tasks as it did not exclusively involve problem-solving but simulated a scavenging situation that involved searching for and obtaining food rewards and allowed for higher options of food sharing. Task 3 further allowed us to investigate social tolerance among group members. Our study was aimed to understand the social tolerance of free-ranging dogs in their natural groups, group task performance and other associated factors like gazing at humans and conspecifics in entirely different contexts. We expected that free-ranging dog groups would perform better in the familiar tasks (Task 2 and 3) than the unfamiliar one (Task 1) and show tolerance among members by sharing abundant resources (Task 3). Based on earlier observations, we also hypothesized that dogs would gaze more towards the human experimenter in the unfamiliar task.

## Materials and Methods

### A. Subjects and Study Sites

The study was carried out in different parts of West Bengal, India. We tested a total of 113 groups of adult free-ranging dogs (summing up to a total of 434 dogs) with group sizes ranging from 3 to 10 (3.65 ± 1.26). Individuals (≥ 3) that were sighted either resting or moving together, with not more than 1 m distance in between, were considered as a group. We used three different tasks for the study. Each group was tested only once with a randomly assigned task. The study was carried out at random locations including residential areas, market places, bus stops, and railway stations between 0900 hours and 1700 hours, during April – July 2016. We carried out the trials in different locations to eliminate the possibility of re-testing a group. Besides, a large area (∼ 456 sq km) was covered to eliminate any re-sampling completely. We relied on the coat colour, scar marks and specific colour patches on the body of the dogs as distinguishing characters for individuals, and the territorial nature of the dogs as identities of the groups tested.

### B. Experimental Procedure

As mentioned above, three different tasks were used in the study, with each group being tested for only one task. For each task, the experimenter (E) walked on random streets in a pre-selected locality in search of groups of free-ranging dogs. On sighting a group, E tried to attract the attention of the individuals by calling out to them prior to the commencement of the trial (see Bhattacharjee et al 2017). All the groups which responded and approached E were used for the task subsequently. Tasks were recorded by a cameraperson from a distance to avoid any interactions with the dogs.

#### Task 1

Free-ranging dogs are accustomed to scavenging from garbage bins, open garbage or closed plastic bags carrying food and/or garbage. This task was designed to mimic a scavenging condition but from an unfamiliar source. It required the dogs to obtain food from a transparent plastic container (0.11 m × 0.11 m × 0.06 m) that had a hole pierced in one corner of the lid through which a nylon rope (length – 0.2 m) had been inserted and attached such that pulling the rope could open the lid of the box. In an earlier experiment, individual free-ranging dogs have been observed to attempt the task but failed to solve it on most occasions (Udell 2015; Brubaker et al. 2017; Bhattacharjee et al. 2017a). Hence this task was considered to be suitable for testing if group members would cooperate to solve the task. E allowed the dogs of the focal group to sniff a boneless raw chicken piece (approximately 0.05 – 0.06 kg in weight) and placed it inside the box. E then placed the box on the ground, approximately 1 m away from the focal group and approximately equidistant to the group members, and moved back to a distance of 0.5 m. Thus, the initial distance between E and the dogs was approximately 1.5 m. E stood in a neutral posture and looked straight ahead without bending his/her head or making eye contact with any of the focal group dogs. The response was recorded for 120 seconds or until the dogs ate the raw chicken piece, whichever was earlier, following which the food was removed. Forty-four adult dog groups were tested for this task.

#### Task 2

In this task, we provided dog groups with a piece of raw chicken as a reward, placed inside a transparent plastic bag (0.19 m × 0.11 m). The experimenter allowed dogs of the focal group to sniff the chicken piece before placing it inside the plastic bag and tying the mouth of the bag with a thread, allowing the dogs to watch the process (Bhattacharjee et al. 2017a). All the other steps were as in Task 1 and the response was recorded for 120 seconds. 43 adult dog groups were tested for this task.

#### Task 3

In this condition, the dog groups were provided with one open plastic basket (0.30 m × 0.07 m × 0.14 m) containing non-edible garbage (dry paper, plastic, leaves etc.) and food rewards, thus emulating a garbage bin (**Fig 1**). The food reward consisted of five pieces of raw chicken and five pieces of bread (representing proteins and carbohydrates respectively), which were mixed with the garbage, as is the case in most waste disposal sites that are accessible to free-ranging dogs in India. Since the task, in this case, did not involve opening the basket to reach the food, the time provided for the task was 60 seconds, instead of 120 seconds, starting after the basket was placed on the ground. All other steps were the same as in the other two tasks. 26 adult dog groups were tested for this task.

**Fig 1.**
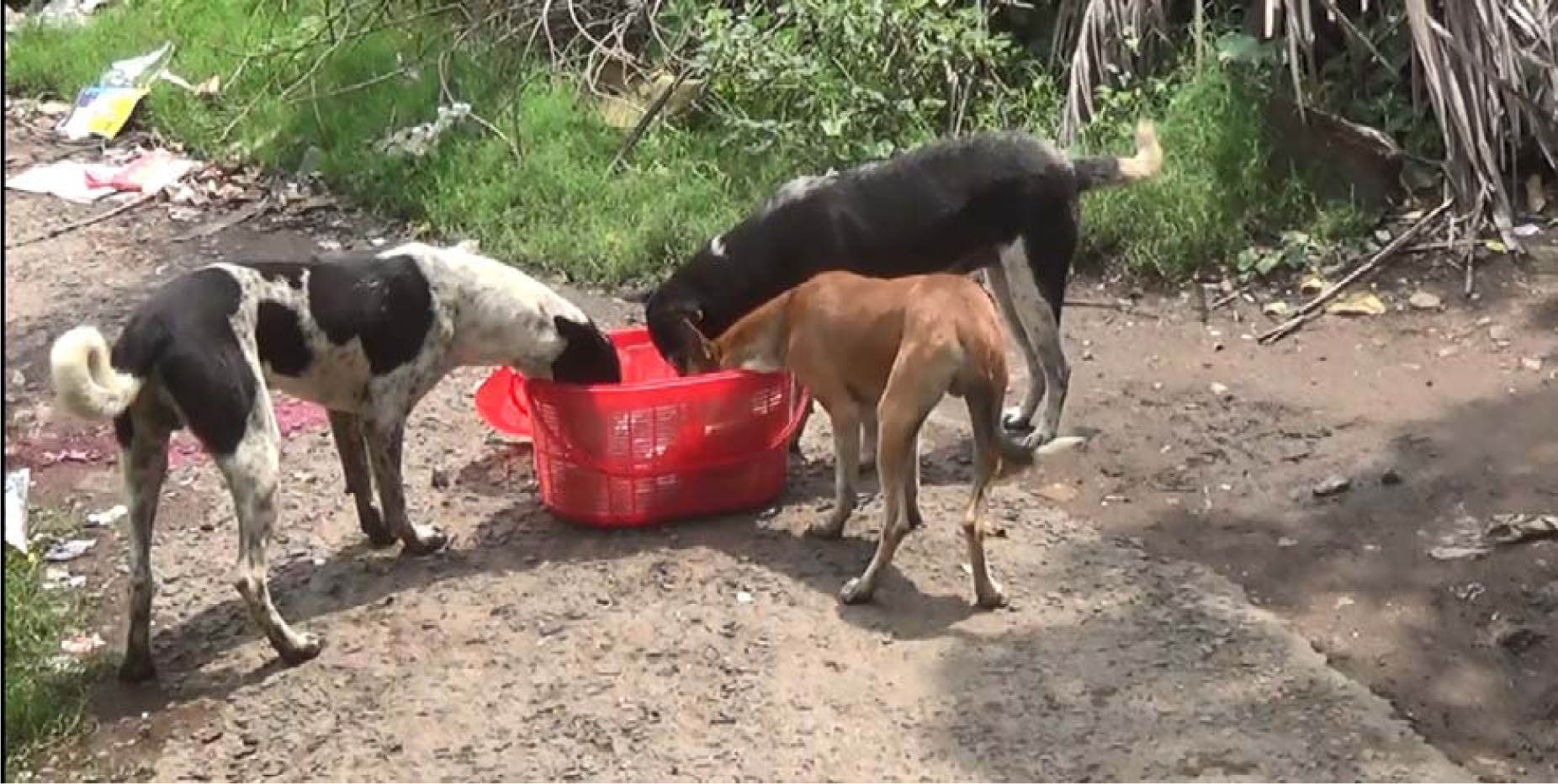
Image showing the experimental set-up of Task 3. Picture courtesy – Shubhra Sau.

The nature of the three tasks differed in terms of their familiarity, quantity of rewards, and to some extent difficulty. Task 1 had earlier been shown to be solved by individual free-ranging dogs, suggesting no physical limitation on part of the dogs. However, a small success rate could be addressed by ‘task difficulty’ along with unfamiliarity, also, Task 2 was highly familiar for these dogs from a scavenging perspective and solved at a higher rate compared to Task 1 (Bhattacharjee et al. 2017a). In order to eliminate any anthropomorphic bias, we have emphasized the familiarity of the tasks (Task 1 and 2), rather than their difficulty levels. Task 3 represented a condition which did not involve problem-solving but allowed us to understand co-feeding and social tolerance. To be better able to understand the various projections of our study, we first compared Task 1 and 2 in order to check for an effect of familiarity and cooperation and later analysed Task 3 (compared a few parameters with tasks 1 and 2) to address whether changes in the quantity of available food resources potentially promotes sharing behaviour/ social tolerance in free-ranging dogs.

### C. Data analysis and statistics

All the videos of task performance and associated behaviours were coded by an individual, which were then used for further analysis. Another individual, blind to the experiment, coded 20% of the data selected randomly. Reliability for success, latency, persistence and gazing measures was found to be high (success: Cohen’s kappa = 0.99; latency: kappa = 0.99; persistence: kappa = 0.96; gazing: kappa = 0.97). Shapiro-Wilk tests were conducted to check for normality of the data. The data were not normally distributed. Thus we performed non-parametric tests. Alpha level was 0.05 throughout the analysis. R Studio and StatistiXL version 1.11.0.0 were used for the analyses.

Following is the list of behaviours/parameters that were quantified from the study –

#### (i) Success

Opening the container or plastic bag and obtaining the food reward was considered as a successful event in tasks 1 and 2 respectively. The success rate in a trial of Task 3 was estimated on the basis of the number of food pieces left after 60 seconds. For example, empty basket (all 5 pieces of bread and 5 pieces of chicken eaten) after a trial corresponds to 100% success. We have analyzed success rates at two different ranges – less than 50% and more than or equal to 50%, in order to get an idea of lower and higher success rates respectively.

#### (ii) Latency

The time between the presentation of the task before the dogs and the display of first response, which involved approach within a distance of 0.05 – 0.1m of the task set-up was defined as latency. We used markers (e.g. leaves, small stones) to get an idea of the distances. Latencies for all the dogs in a group were recorded but only the latency of the first dog that approached a task was considered for the analysis.

#### (iii) Persistence

Persistence was defined as the duration of active engagement or involvement in the task. We considered active engagement when dogs showed the following behaviours with the objects (box/bag/basket) - ‘touching’, ‘licking’, ‘pulling’, and ‘obtaining the food reward(s)’. Persistence was exclusive of the duration of interruptions when dogs were not actively engaged in task solving. Calculation of persistence was cumulative. We quantified (a) Persistence of individuals (persistence of each group member), (b) Group persistence (average persistence of the members of a group) and (c) Persistence of the solving individual (persistence of a group member that finally solved a task, Task 1 and 2 specific).

#### (iv) Cooperation (Task 1 and 2 specific)

Two or more individuals of a group acting together, without aggressive interactions, to solve a task was considered as cooperation or simultaneous engagement at the task. We calculated the number of individuals (at least 2) persisting on a task and the duration of overlap to define cooperation.

#### (v) Social tolerance (Task 3 specific)

Social tolerance was defined as a tendency of the group members to scavenge from the same resource side by side without aggression. In order to measure this, a ‘Tolerance Index’ (ToI) was constructed for each individual in a group. ToI intended to evaluate the extent to which the group members performed the task together and was not meant to compute the evolutionary benefits being incurred by the individuals due to such an action.

We used the following parameters while constructing ToI:

- Number of individuals that a focal dog can interact with - for example, in a group of 4 individuals, a focal dog would be able to interact with a maximum of 3 individuals.
- Availability of time to solve a task - here we subtracted the latency from the total task duration. For example, in a task of 120 seconds, a focal dog with a latency of 10 seconds would have 110 seconds of time available for cooperation.
- Overlap with other members - we calculated the number of individuals that were already engaged in the task when a focal dog joined. Similarly, the duration of the overlap was also calculated. For example, in a group of 4 individuals, a focal dog’s active engagement with a task overlapped with 2 other members of the group for 30 seconds and with another member for 10 seconds. While calculating ToI, we first multiplied the proportion of individuals that the focal dog tolerated (2/3 and 1/3) with the proportion of time available that it spent with each in the task (30/110 and 10/110, considering the availability of task time as 110 sec), and then added the two values {(2/3*30/110) + (1/3*10/110)}.
- Leaving – We used two parameters to assess the situation at the point when a focal dog left the task; the time remaining for the task and the proportion of group members engaged in the task. For example, if the above focal dog left the task at 90 seconds while two of the group members were actively engaged with the task at that time, this factor was calculated as 2/3*20/110.

We used the following formula to calculate ToI –

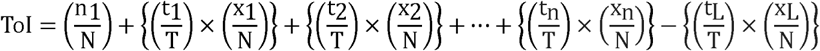

[N = (Total group size – 1); i.e., the number of individuals in the group a focal dog can interact with; T = Total duration of the experiment - latency of the focal animal; n_1_ = number of individuals engaged in the task when the focal dog joins; t_n_ = duration of overlap with x_n_ number of individuals; t_L_ = time remaining for the experiment when the focal dog leaves the task; x_L_ = number of individuals engaged in the task when the focal dog leaves].

We calculated the ToI values for the individuals that approached in Task 3. Lesser ToI value of an individual indicated a lower tendency to act together with its group members, i.e., a lower intention for food sharing and cooperation. We also calculated the mean ToI values of the groups to check for any correlation with corresponding success rates.

#### (vi) Food sharing

Sharing of food rewards without aggression among the group members (at least within 2 members) was considered as food sharing. For task 3, co-feeding was the proxy for food sharing. Co-feeding was determined by calculating the percentage of group members feeding together in Task 3. For example, in a group of 4 individuals, 100% sharing indicated that all the group members had fed/scavenged together, whereas, 75% sharing was recorded when 3 of them was observed to co-feed.

#### (vii) Gazing

The duration of gazing at the upper body of the human experimenter was recorded. Gazing towards the conspecifics was also quantified.

#### (viii) Aggression

Aggressive behaviours were aimed towards the conspecifics and included threatening responses. We quantified the following behaviours as aggressive during the tasks: snarling (aggressive vocal response to a group member), threatening (growling/barking at another dog with alert posture having ears pointed) and biting. Neutral and affiliative responses were treated as no aggression. Affiliative behaviours included proximity seeking, contact seeking, social facilitation, tail wagging, and relaxed posture, while neutral responses were restricted to resting, self-care (scratching, licking, grooming) and general disinterest.

#### (ix) First inspection, highest persistence and retrieval of food reward

Since free-ranging dogs are scavengers, we hypothesize that an opportunistic individual would inspect a task first, persist most and obtain the food in case of Task 1 and 2, illustrating a strategy of 1-1-1 (rank 1 for inspection, persistence and retrieval of food reward). For Task 3 it was difficult to gauge the actual amount of food obtained by an individual but we assumed the time spent by an individual in feeding as a correlate of the amount of food eaten. Groups that failed to obtain food rewards were not considered for this calculation.

## Results

### Task 1 vs Task 2

#### (i) Success

The dog groups performed significantly better in Task 2 than in Task 1 **(Fig. 2)**. Success rates for Task 2 and 1 were 95% and 23% respectively (Chi-squared goodness of fit, χ ^2^ = 45.547, d_Cohen_ = 2.096, N = 87, df = 1, p < 0.0001).

**Fig 2.**
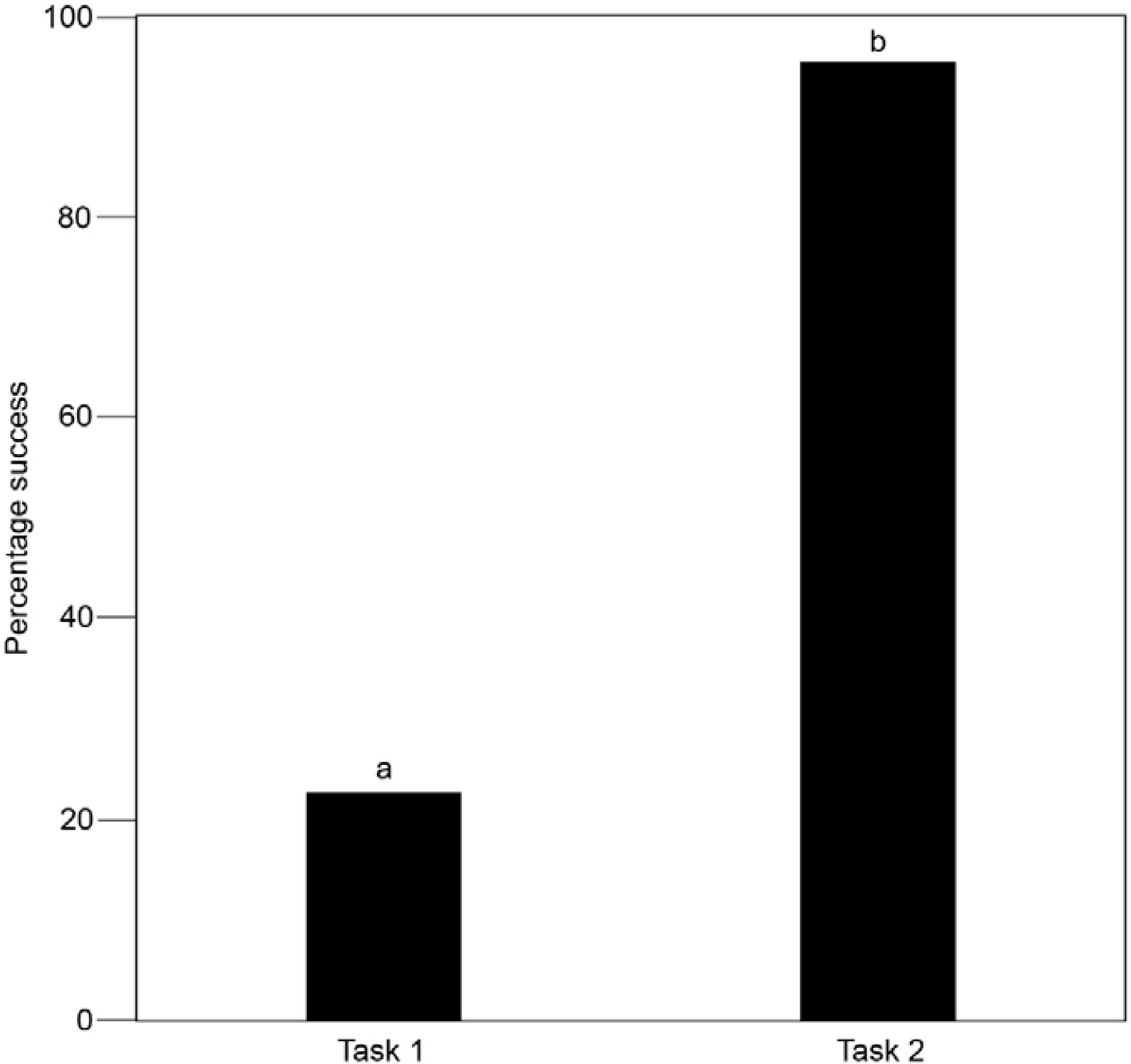
Success rates in task 1 (unfamiliar) and task 2 (familiar). Bar graph showing percentage of groups that successfully solved the two tasks. Groups in task 2 showed significantly higher success rates that task 1. Different letters indicate a significant difference between the categories.

#### (ii) Latency

Latencies varied significantly between Tasks 1 and 2 (Mann-Whitney U test, U = 1160.500, d_Cohen_ = 0.45, N = 86, df1 = 43, df2 = 43, p = 0.04). Individuals from one group did not respond in Task 1 and hence the total sample size reduced by 1 to 86. Dogs showed significantly faster response (1.16 ± 0.37 sec) in Task 2 as compared to Task 1 (1.60 ± 0.90 sec).

#### (iii) Persistence

(a) Persistence of individuals - members of a group that approached a task were considered for the analyses (Sample size: Task 1 – 126, Task 2 – 92). We obtained no difference between Tasks 1 and 2 (Mann-Whitney U test, U = 6288.000, d_Cohen_ = 0.145, df1 = 126, df2 = 92, p = 0.286). (b) Group persistence – Average persistence of the groups did not differ between Tasks 1 and 2 (Mann-Whitney U test, U = 997.000, d_Cohen_ = 0.093, df1 = 44, df2 = 43, p = 0.670). (c) Persistence of the solving individuals - There was no significant difference in persistence between the individuals that finally solved Tasks 1 and 2 (Mann-Whitney U test, U = 218.000, d_Cohen_ = 0.086, df1 = 10, df2 = 41, p = 0.770).

#### (iv) Cooperation

We found a difference in the duration of cooperation between the tasks (Mann-Whitney U test, U = 1813, d_Cohen_ = 0.793, df1= 54, df2 = 47, p < 0.0001). Groups in Task 1 engaged with the task together longer (4.94 ± 6.37 sec) compared to Task 2 (1.08 ± 1.62). In both the tasks, ‘pairs’ from the groups were seen as cooperating units more often than ‘triads’ and ‘tetrad or more’ (Task 1 - Kruskal-Wallis test, χ^2^ = 15.648, d_Cohen_ = 1.209, df = 2, p < 0.0001, Task 2 - Kruskal-Wallis test, χ^2^ = 28.224, d_Cohen_ = 2.429, df = 2, p < 0.0001, **Fig 3, Supplementary Text 1**).

**Fig 3.**
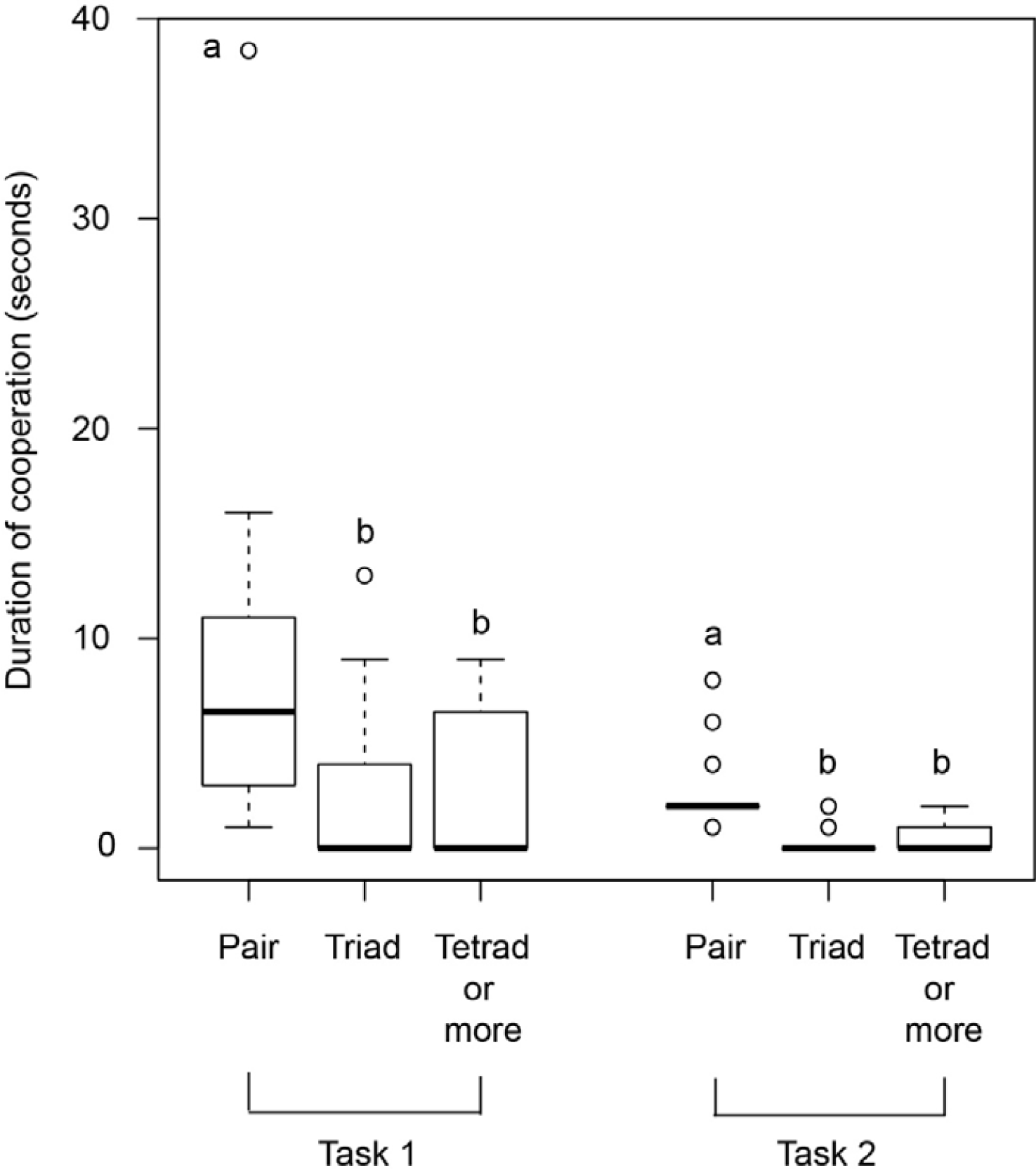
Duration of cooperative task solving of groups (Task 1 and 2) in ‘pair’, ‘triad’ and ‘tetrad or more’ formations. Box and whisker plot illustrating the duration of group task solving in Task 1 and 2. Both tasks 1 and 2 showed that the ‘pairs’ from the groups worked longer as cooperating units than ‘triads’ and ‘tetrad or more’ (Task 1 - Kruskal-Wallis test, χ^2^ = 15.648, d_Cohen_ = 1.209, df = 2, p < 0.0001; Task 2 - Task 2 - Kruskal-Wallis test, χ^2^ = 28.224, d_Cohen_ = 2.429, df = 2, p < 0.0001). Boxes represent interquartile range, horizontal bars within boxes indicate median values, and whiskers represent the upper range of the data. Different letters indicate a significant difference within tasks.

#### (v) Food sharing

We found absolutely zero sharing of food in Tasks 1 and 2. None of the group members shared the retrieved food rewards with their conspecifics.

#### (vi) Gazing

Dogs gazed at the human experimenter significantly more than the conspecifics in Task 1 (Mann-Whitney U test, U = 29424.500, d_Cohen_ = 2.639, df1 = 175, df2 = 175, p < 0.0001). However, in Task 2, there was no such difference (Mann-Whitney U test, U = 13675.500, d_Cohen_ = 0.118, df1 = 160, df2 = 160, p < 0.291).

#### (vii) Aggression

Dogs displayed very less aggression towards their group members in both the tasks. Approximately 7% and 14% of the dogs showed aggressive behaviours in Tasks 1 and 2 respectively; the level of aggression was comparable between the two tasks (Chi-squared goodness of fit, χ^2^ = 2.333, d_Cohen_ = 0.332, N = 87, df = 1, p = 0.12). Aggression was less compared to all the other (both affiliative and neutral together) behaviours displayed during the tasks (Task 1: Goodness of fit, χ ^2^ = 32.818, d_Cohen_ = 3.426, N = 44, df = 1, p < 0.0001; Task 2: Goodness of fit, χ^2^ = 22.349, d_Cohen_ = 2.080, N = 43, df = 1, p < 0.0001).

#### (viii) First inspection, highest persistence and retrieval of food reward

A total of 51 groups of dogs successfully solved Tasks 1 and 2 (Task 1 –10, Task 2 – 41). We pooled data from both the tasks to estimate the proportion of groups in which the first individual to respond to the task was also the one to have persisted the longest and solved the task. In 37 out of 51 groups, the individual which inspected a task first showed highest persistence and also retrieved the reward. We found a difference between the groups that showed a first inspection – highest persistence – retrieval of food reward strategy and groups that did not (Goodness of fit, χ^2^ = 10.373, d_Cohen_ = 1.01, N = 51, df = 1, p = 0.001).

### Task 3

#### (i) Success

Out of 26 groups, only 3 groups showed 100% success and 2 groups showed zero success. However, we found no difference between the two ranges of the success rates considered for Task 3 (lower (< 50%) and higher - (≥ 50%) success rates; Chi-squared goodness of fit, χ^2^ = 1.385, d_Cohen_ = 0.474, N = 26, df = 1, p = 0.239), indicating a somewhat uniform distribution between 0% - 100% **(Fig 4)**.

**Fig 4.**
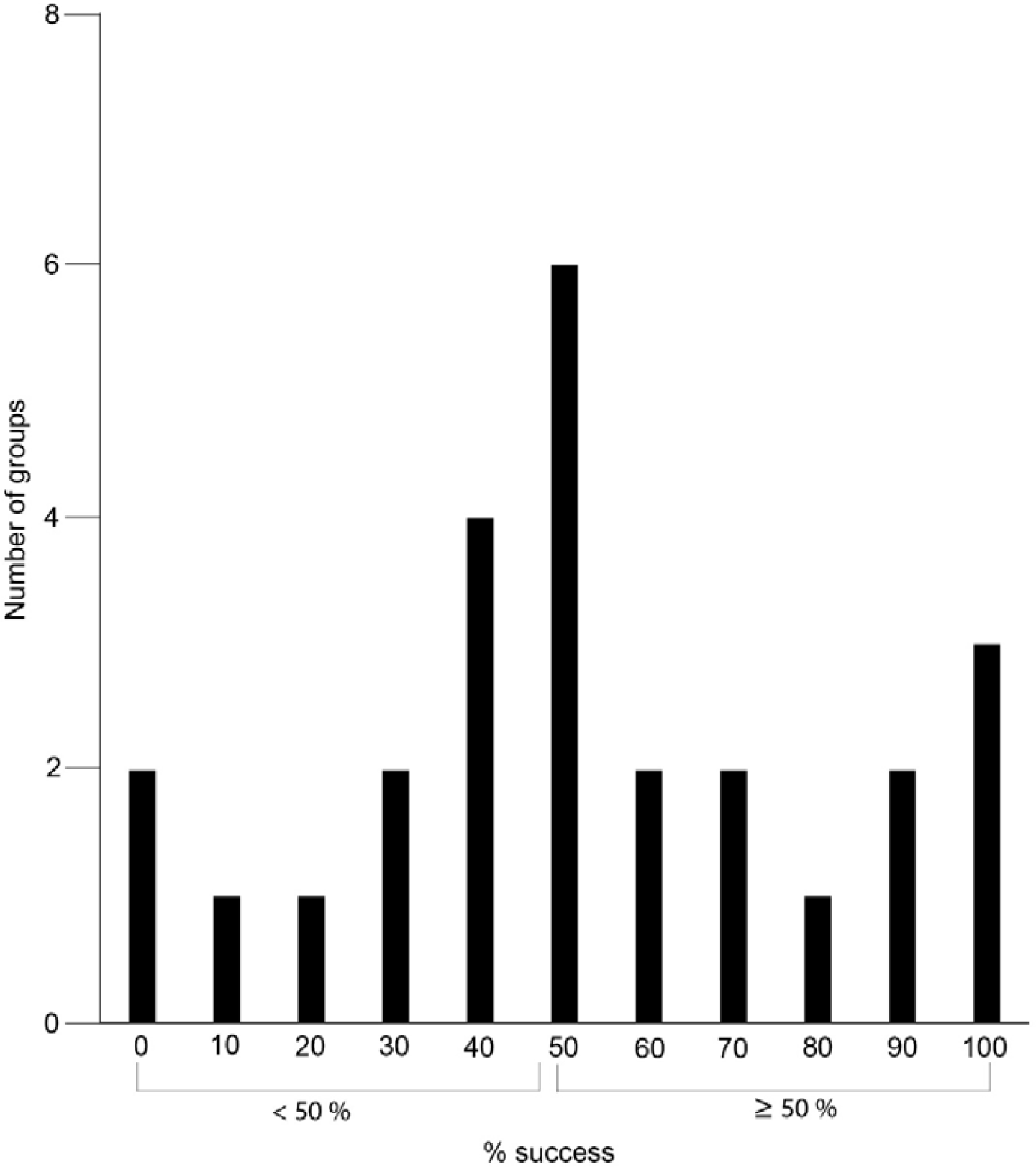
Success rates in task 3. Histogram showing the success rates in corresponding number of groups in Task 3. No difference was found between lower success (< 50%) and higher success (≥ 50%) rates (Chi-squared goodness of fit, χ^2^ = 1.385, d_Cohen_ = 0.474, N = 26, df = 1, p = 0.239).

#### (ii) Latency

Dogs appeared to be quite hesitant in approaching Task 3 (6.84±7.26 sec). Latency of the dogs in Task 3 differed from both Task 1 (Mann-Whitney U test, U = 881.000, d_Cohen_ = 1.094, df1 = 43, df2 = 26, p < 0.001) and 2 (Mann-Whitney U test, U = 972.000, d_Cohen_ = 1.563, df1 = 43, df2 = 26, p < 0.001).

#### (iii) Persistence

(a) Persistence of individuals – A total of 58 individuals, considering all the groups, persisted in Task 3. Persistence of individuals was found to be higher in Task 3 compared to individuals in Task 1 (Mann-Whitney U test, U = 5875.500, d_Cohen_ = 1.118, df1 = 126, df2 = 58, p < 0.001) and Task 2 (Mann-Whitney U test, U = 4184.500, d_Cohen_ = 1.088, df1 = 92, df2 = 58, p < 0.001).

(b) Group persistence – We found the average group persistence between the three tasks to be significantly different (Kruskal-Wallis test, χ^2^ = 27.053, d_Cohen_ = 1.086, df = 2, p < 0.0001). Post-hoc pairwise comparisons revealed similar outcomes as found in individual persistence. In Task 3, groups showed higher persistence compared to Task 1 (Mann-Whitney U test, U = 962.000, d_Cohen_ = 1.375, df1 = 44, df2 = 26, p < 0.0001) and Task 2 (Mann-Whitney U test, U = 929.500, d_Cohen_ = 1.325, df1 = 43, df2 = 26, p < 0.0001).

#### (iv) Social tolerance

ToI values of the all the dogs ranged between 0 and 1 (0.22 ± 0.31). We found a strong positive correlation between food sharing in the groups and their ToI values (Spearman rank correlation, r_s_ = 0.816, df = 26, p < 0.001, **Fig 5**). However, there was no correlation between success rates (feeding) of the groups and their ToI values (Spearman rank correlation, r_s_ = 0.281, df = 26, p = 0.164).

**Fig 5.**
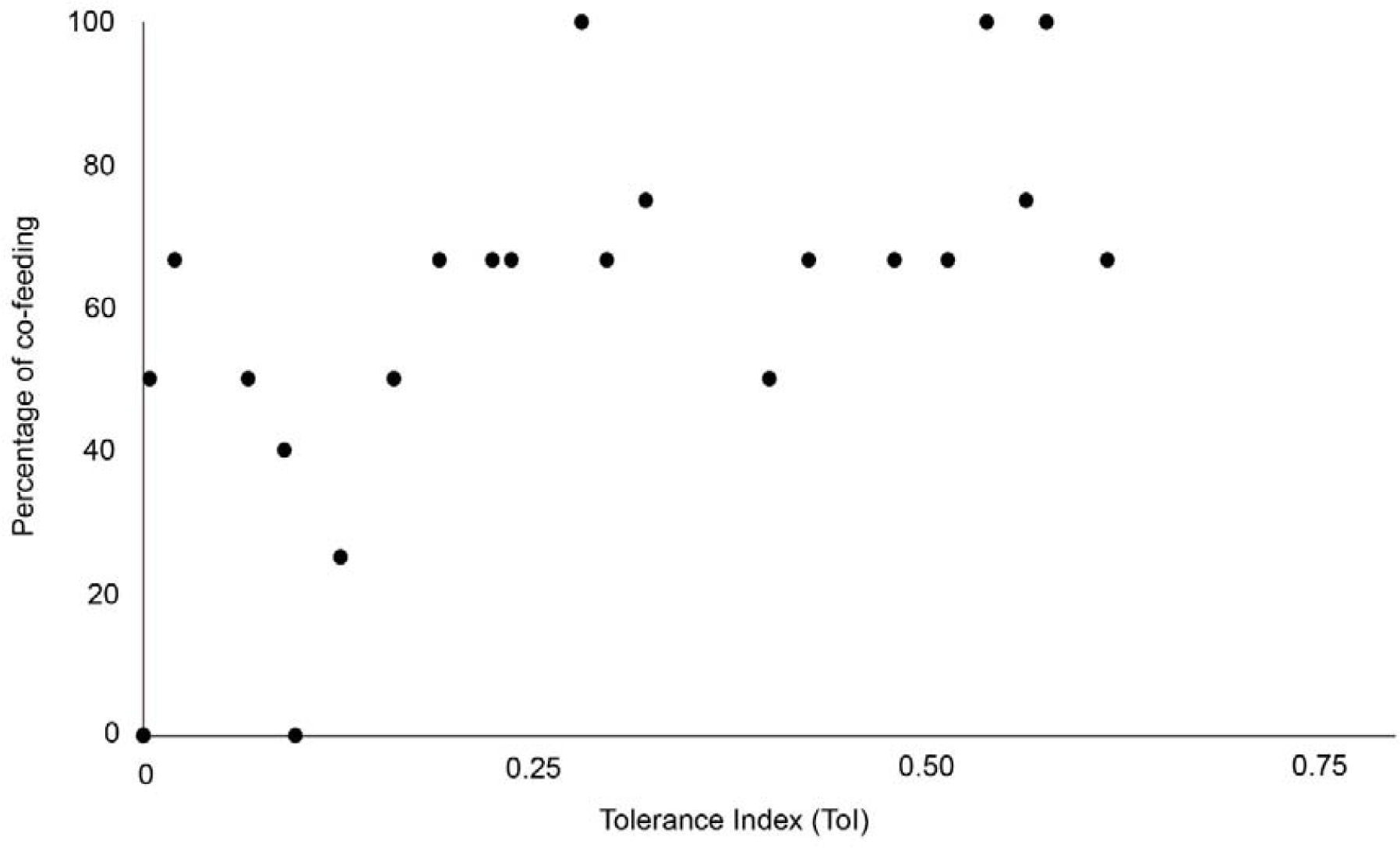
Correlation between ToI and food sharing in Task 3. Spearman-rank correlation showed a strong positive relationship between average ToI and food sharing in groups of Task 3 (r_s_ = 0.816). Group food sharing values (in percentage) were plotted on the Y-axis and ToI values on the X-axis.

#### (v) Gazing

Dogs displayed higher gazing at the conspecifics than the human experimenter in Task 3 (Mann-Whitney U test, U = 6080.000, d_Cohen_ = 0.425, df1 = 99, df2 = 99, p = 0.003).

#### (vi) Aggression

Close to 11% of the dogs elicited aggressive responses towards their group members. Aggression was significantly less compared to neutral and affiliative behaviours together (Goodness of fit, χ^2^ = 15.385, d_Cohen_ = 2.407, N = 26, df = 1, p < 0.0001).

#### (vii) First inspection, highest persistence and retrieval of food reward

We considered 22 groups that responded and persisted in Task 3 for this calculation. Similar to Task 1 and 2, we found groups that showed the 1-1-1 condition higher in Task 3 (Goodness of fit, χ ^2^ = 8.909, d_Cohen_ = 1.649, N = 22, df = 1, p = 0.003).

## Discussion

Our study revealed that free-ranging dog groups performed better in the familiar task (Task 2) as compared to the unfamiliar one (Task 1) when faced with tasks with single food rewards. This was consistent with the earlier findings with individual free-ranging dogs and further substantiates dogs’ inferior abilities in physical cognitive task solving situations like string pulling (Osthaus et al. 2005; Bhattacharjee et al. 2017a). Dogs also showed a much faster reaction to Task 2 than Task 1, emphasizing the role of familiarity (Bhattacharjee et al. 2017a). In case of Task 3, the uniform distribution of success rates across 0 – 100% did not help in providing any useful insights into the free-ranging dogs’ scavenging abilities. In spite of its familiar nature, dogs took longer to approach the set-up in Task 3 relative to the other tasks. Such outcomes could be attributed to fear or hesitation due to the unusual way of food provisioning by an unknown human. Free-ranging dogs in India are generally reluctant to approach garbage bins while humans are still around for disposal of garbage/leftover food as, in such cases, dogs are, typically, shooed away, threatened or beaten by people. A very recent study quantifying free-ranging dog’s scavenging efficiency reported similar results (Sarkar et al. 2019) Thus, we suspect the reasons mentioned above to be the cause for longer approach time of dogs in Task 3.

Dogs displayed higher cooperation in Task 1 compared to Task 2, which could be attributed to an unfamiliar nature and difficulty level of the task. However, success did not depend on cooperation in those tasks. We also reckon two underlying factors that could have influenced the outcomes - (i) non-availability of a set-up where group members of a large group can perform together, and (ii) presence of a moderately large single food reward. It was also noted that members who inspected a task first, persisted more and obtained the food reward and no food sharing was observed in any of the two tasks. This is suggestive of the presence of a feeding hierarchy in free-ranging dog groups, as aggression over the reward was not observed. It is difficult to examine and disentangle the factors mentioned above with the current experimental set-up and require further experimentation. The shortcomings were overcome to some extent with the experimental design of Task 3. It provided evidence of social tolerance, co-feeding and thereby general cooperation among the group members when higher food rewards were available. The minimal display of aggressive behaviour towards each other also corroborated the existence of some level of understanding of hierarchy within the group. Both the individual and group average persistence were found to be higher in Task 3, which could be attributed to the lower effort required to find food and the higher quantity of food available. In a nutshell, differing food levels affected the group responses strikingly.

Gazing responses provided significant information relating to both intra and interspecific communication. Gazing has been considered as a striking means of communication in canids (Miklósi et al. 2000; Hare and Tomasello 2005). It has previously been shown that dogs gaze at human experimenters when faced with unfamiliar or difficult tasks (Szetei et al. 2003; Udell 2015; Brubaker et al. 2017; Bhattacharjee et al. 2017a). The higher rates of gazing at the human experimenter in Task 1 could be associated with information seeking. This statement is further strengthened by the lack of gazing responses during Task 2, when the dogs faced a familiar task that they could solve independently (Bhattacharjee et al. 2017a). However, this does not rule out the chances of dogs being vigilant or alert when encountering an unfamiliar human at proximity. Relatively higher gazing at the group members during Task 3 is likely to be used for figuring out other group members’ intentions to approach the task. Studies have concluded that being able to estimate the intentions of others is vital across social contexts like maintenance of social cohesiveness (Cheney et al. 1986; Friedkin 2004), availing sneaky mating opportunities (Alberts et al. 2003), territorial defence (Crockford et al. 2012), challenging the higher ranking individuals to take over a group (Rowell 1974) etc. It might also help dogs to maintain cohesion and structure within the group during scavenging.

Members of free-ranging dog groups seemed to lack the tendency to perform together in solving physical cognitive tasks, but they do show tolerance towards each other, resulting in considerably higher food sharing. Cooperation in the form of social tolerance and co-feeding is a primer for a higher level of cognitive complexity, which requires coordination and communication between group members. The current set of experiments demonstrate that free-ranging dog groups are capable of showing social tolerance towards group members during scavenging and can also exhibit co-feeding, but further experimentation is required to investigate how the groups are maintained and persist over time. From these experiments, we conclude that free-ranging dogs can cooperate with their group members during scavenging, but choose to do so based on context, e.g. when high rewards are available. This observation substantiates earlier observations that free-ranging dogs tend to scavenge solitarily most of the time, possibly to avoid potential conflict, but forage in pairs or larger groups in social contexts like mating and pup rearing (Sen Majumder et al., 2014). This study provides further testimony to the flexible social organization in these dogs, which demonstrate interesting cooperation-conflict dynamics within their social groups. Free-ranging dogs survive in a human-dominated environment, where aggression between dogs is met with intolerance from humans. Hence, maintaining feeding hierarchies using aggression or fighting over resources is not adaptive for the dogs. On the other hand, the use of subtle cues like display of dominance and subordination through postures and vocalizations might be an effective mechanism for maintaining hierarchies that help to develop efficient scavenging strategies. Long term observations of group-level behaviour of the free-ranging dogs would help to provide deeper insights into how such hierarchies are established and maintained.

## Acknowledgements

We would like to thank Ms. Ankurita Mondal for helping with the video recording for some of the trials.

## Funding

This study was partially supported by the SERB Women’s Excellence Award to AB (SB/WEA-005/2013). DB was supported by a DST INSPIRE Fellowship. AA was supported by the IASc-INSA-NASI Summer Research

Fellowship program; DM was supported by IISER Kolkata summer fellowship program. We thank Indian Institute of Science Education and Research (IISER) Kolkata for providing infrastructural support for this work.

## Data availability

The datasets analysed during the current study are available from the corresponding author on reasonable request.

## Compliance with ethical standards

### Conflict of interest

The authors declare that they have no conflict of interest.

### Ethical approval

The study design did not violate the Animal Ethics regulations of the Government of India (Prevention of Cruelty to Animals Act 1960, Amendment 1982). The protocol for the experiment was approved by the IISER Kolkata Animal Ethics Committee, as part of a larger project sanctioned by the SERB (EMR/2016/000595).

### Informed consent

All the human participants involved in this study gave their consent.

